# Modelling somatic mutation accumulation and expansion in a long-lived tree with hierarchical modular architecture

**DOI:** 10.1101/2022.05.02.490262

**Authors:** Sou Tomimoto, Akiko Satake

## Abstract

In a long-lived organism with a modular architecture, such as trees, somatic mutations accumulate throughout the long lifespan and result in genetic mosaicism in each module within the same individual. In recent years, next-generation sequencing technology has provided a snapshot of such intra-organismal genetic variability. However, the dynamic processes underlying the accumulation and expansion of somatic mutations during the growth remain poorly understood. In this study, we constructed a model to describe these processes in a form that can be applied to a real tree. Given that the proliferation dynamics of meristematic cells vary across plant species, multiple possible processes for elongation and branching were comprehensively expressed in our model. Using published data from a poplar tree, we compared the prediction of the models with the observation and explained the cell lineage dynamics underlying somatic mutations accumulation that were not evident from the snapshot of the sequenced data. We showed that the somatic genetic drift during growth increases inter-meristem mosaicism, resulting in genetically distinct branches and less integrity within an individual tree. We also showed that the somatic genetic drift during branching leads to the mutation accumulation pattern that does not reflect the tree topology. Our modelling framework can help interpret and provide further insights into the empirical findings of genetic mosaicism in long-lived trees.

## 1. Introduction

In long-lived organisms, somatic mutations accumulate during mitotic growth and tissue regeneration because the DNA of somatic cells is continuously exposed to endogenous (e.g., production of reactive oxygen species) or exogenous damage (e.g., ionizing radiation and ultraviolet light) and replication errors. In unitary animals, mutations within somatic cells are restricted to the individual in whom they occur because only mutations in the germline are passed on to further generations. In contrast, in plants, there is no distinction between the germline and the soma, leading to the transmission of somatic mutations to progeny (Whitham and Slobodchikoff, 1981; Sutherland and Watkinson, 1986). Therefore, somatic mutations can contribute to the genetic diversity and evolutionary rates within a plant population.

The proliferation and fixation dynamics of *de novo* somatic mutations within individuals and their contradictory roles in individual fitness, mutational meltdown or promotion of adaptation to changing environments, have been discussed since the 1980s (Gill et al., 1995; Klekowski, 2003). With the recent revolution and advance in sequencing technology, it is now possible to measure the quantity and distribution of somatic mutations in the modular architecture of tree individuals. In a 234-year-old oak tree, the number of somatic mutations has been reported to be surprisingly low compared to the value expected from the mutation rate in an annual herb, *Arabidopsis thaliana* (Schmid-Siegert et al., 2017). Similarly, recent studies reported low mutation rates in *Quercus robur* (Plomion et al., 2018), *Picea sitchensis* (Hanlon et al., 2019), *Eucaliptis melliodora* (Orr et al., 2020), and *Populus trichocarpa* (Hofmeister et al., 2020). These somatic mutations were distributed along with the modular architecture of trees (Schmid-Siegert et al., 2017). In addition to genetic variations among somatic cells, somatic epigenetic variations were also detected in *P.trichocarpa* (Hofmeister et al., 2020).

These new empirical findings can largely benefit from mathematical models that integrate the dynamic processes of tree growth and mutation accumulation. Klekowski and Kazarinova-Fukshansky (1984a) developed a model that describes the frequency and fixation dynamics of the neutral somatic mutations during growth. The model was extended to consider the selection of somatic mutations during tree growth (Klekowski and Kazarinova-Fukshansky, 1984b; Pineda-Krch and Fagerström, 1999; Otto and Orive, 1995; Orive, 2001), stratified structure of the meristem (Klekowski et al., 1985; Pineda-Krch and Lehtilä, 2002), modular architecture of the trees (Klekowski et al., 1989), and competition among buds (Antolin and Strobeck, 1985; see Folse and Roughgarden, 2012 for review of the models). Despite the wealth of theoretical studies, no study has developed a model at the whole tree level that can be directly comparable to empirical data.

Here, we modelled cell lineage dynamics in the shoot meristem and predicted accumulation and expansion patterns of somatic mutations in a form that can be applied to a real tree with hierarchical modular architecture. Considering both elongation and branching processes, we evaluated the effects of somatic genetic drift during each process and compared the model predictions with the sequenced data of a poplar tree (Hofmeister et al., 2020). Our approach will be suitable for the genomic revolution era where somatic mutations can be detected quantitatively in diverse organisms with modular architecture.

## 2. Two major growth processes that generate the modular architecture of trees

### 2.1. Elongation: different organization and proliferation dynamics of meristematic cells across plant taxa

The growth of the above-ground tissue of plants results from successive divisions of a collection of stem cells positioned at the summit of the shoot meristem. Here, we call this process “elongation” (Fig. 1). During elongation, stem cells in the shoot meristem maintain the capacity of proliferation.

**Figure 1.**
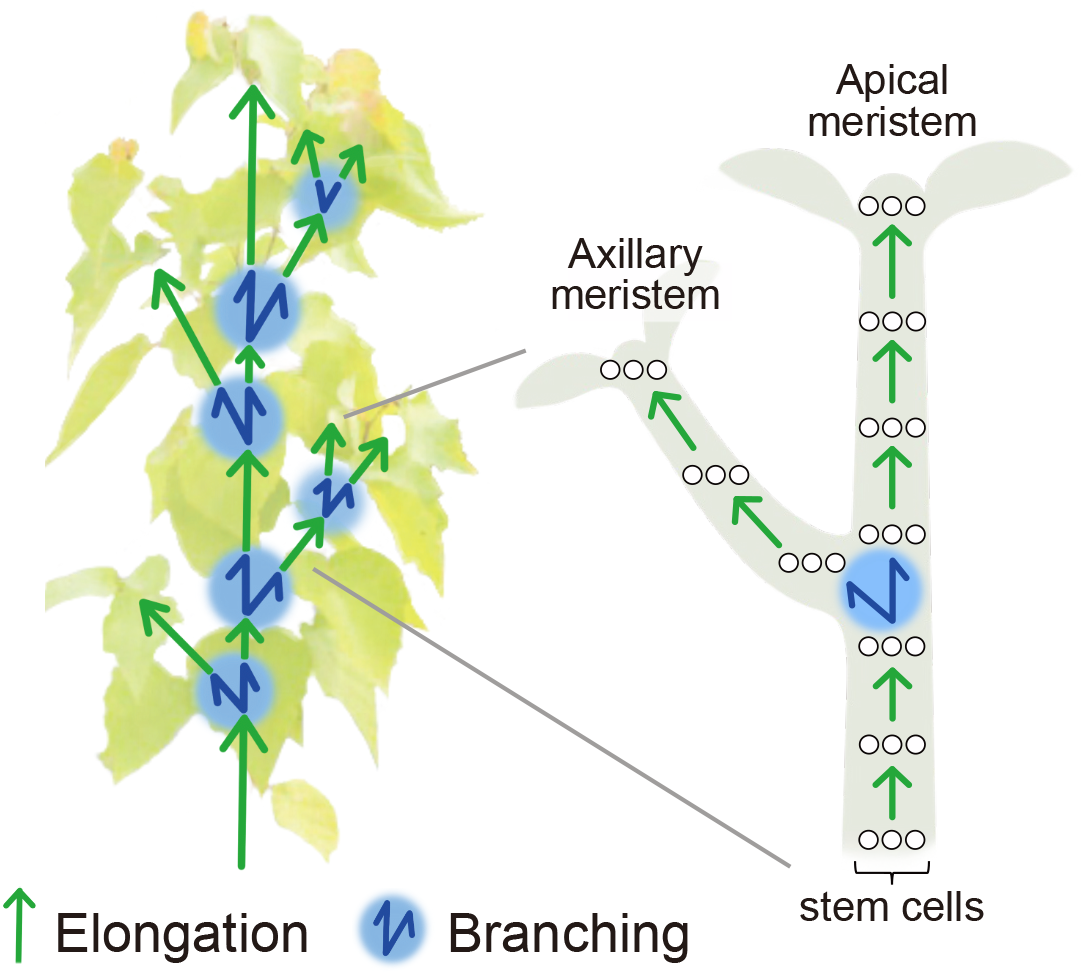
Formation of the modular architecture of the tree as a combination of elongation and branching. One direction arrow (green) and two-way direction arrow (blue) indicate elongation and branching, respectively. Stem cell dynamics along with elongation and branching are illustrated for one of the branches, where a white circle indicates stem cells in the shoot meristem (The number of stem cells *α* = 3).

The number and organization of stem cells in the shoot meristem are different among taxa (Dermen, 1969). In bryophytes and pterophytes, a shoot meristem consists of a single stem cell. In this case, divisions of the single stem cell give rise to all above ground tissue of the individuals (Klekowski, 1984). On the other hand, the meristem of seed plants consists of multiple stem cells and possesses a more complex layered organization called the tunica-corpus structure (Schmidt, 1924). During elongation, stem cells in the outer tunica layers divide anticlinally, and an inner corpus layer divides both anticlinally and periclinally (Satina, et al., 1940; Poething, 1989; Frank and Chitowood, 2016). Occasionally, however, periclinal division of tunica cells occurs and leads to the migration of cell lines from the tunica into the corpus. This process results in the instability of stem cell lineages (Dermen, 1969; Zahrdadníková, et al., 2020). The stability of the tunica-corpus cell lineage differes across species (Poething, 1989). The analysis of plant chimaeras showed that the most angiosperms have stable tunica-corpus distinctions in which the cell lineages derived from a specific layer are usually bound to that layer due to strict anticlinal division. In contrast, gymnosperm species are more likely to divide periclinally (Dermen, 1969). This process leads to less pronounced tunica-corpus distinction and displacement of cells by the other, in most cases invasion of tunica cells into corpus, resulting in the lack of permanent stem cell initials (Poething, 1989; Klekowski, 2003; Zahrdadnikova et al., 2020). Given that the difference in the stability of the tunica-corpus distinction can influence the accumulation and proliferation patterns of somatic mutations, we developed models for different types of meristems.

### 2.2. Branching: axillary meristem formation and sampling of cell lineages

Another growth process involved in the generation of the modular architecture of trees is the formation of an axillary meristem from the apical meristem via a process called “branching” (Fig. 1). An axillary meristem gives origin to a lateral branch. Along with the successive divisions of stem cell initials, cells proliferating from the initials expand radially (Marc and Hackett, 1992; Hara, 1995) and form a new axillary meristem on the apical meristem flanks (Burian et al., 2016). Thus, not all stem cell lineages, but rather only a subset of them, contribute to the formation of an axillary meristem. This process may lead to the biased sampling of cell lineages. Given the difference in the process of axillary meristem formation, the degree of bias during the sampling of stem cells may vary across species (Dermen, 1969; Zahrdadníková, et al., 2020).

## 3. Model

The modular architecture of trees is formed through the reiteration of elongation and branching (Fig. 1; Antolin and Strobeck, 1985; Godin and Caraglio, 1998). To describe the behaviour of somatic mutations at the whole tree level, we modelled stem cell population dynamics during both the elongation and branching processes. For simplicity, we assumed that the number of stem cells remains constant, *α*, throughout the growth in both the apical and axillary meristems.

### 3.1. Modelling elongation

Shoot elongation in seed plants can generally be modelled for two different types of meristems, structured or stochastic meristems (Klekowski and Kazarinova-Fukshansky, 1984a). Structured meristems are characterized by the strict tunica-corpus distinction frequently observed in angiosperms, and the lineages of cells that arise from a specific layer are relatively permanent resident initial stem cells. This structured meristem is modelled by assuming that one of the daughter cells of each stem cell is persistent after cell division and that the remaining daughter cells differentiate into different cell types (Fig. 2a). In other words, stem cell lineages in the meristem of the structured model are maintained during elongation.

**Figure 2.**
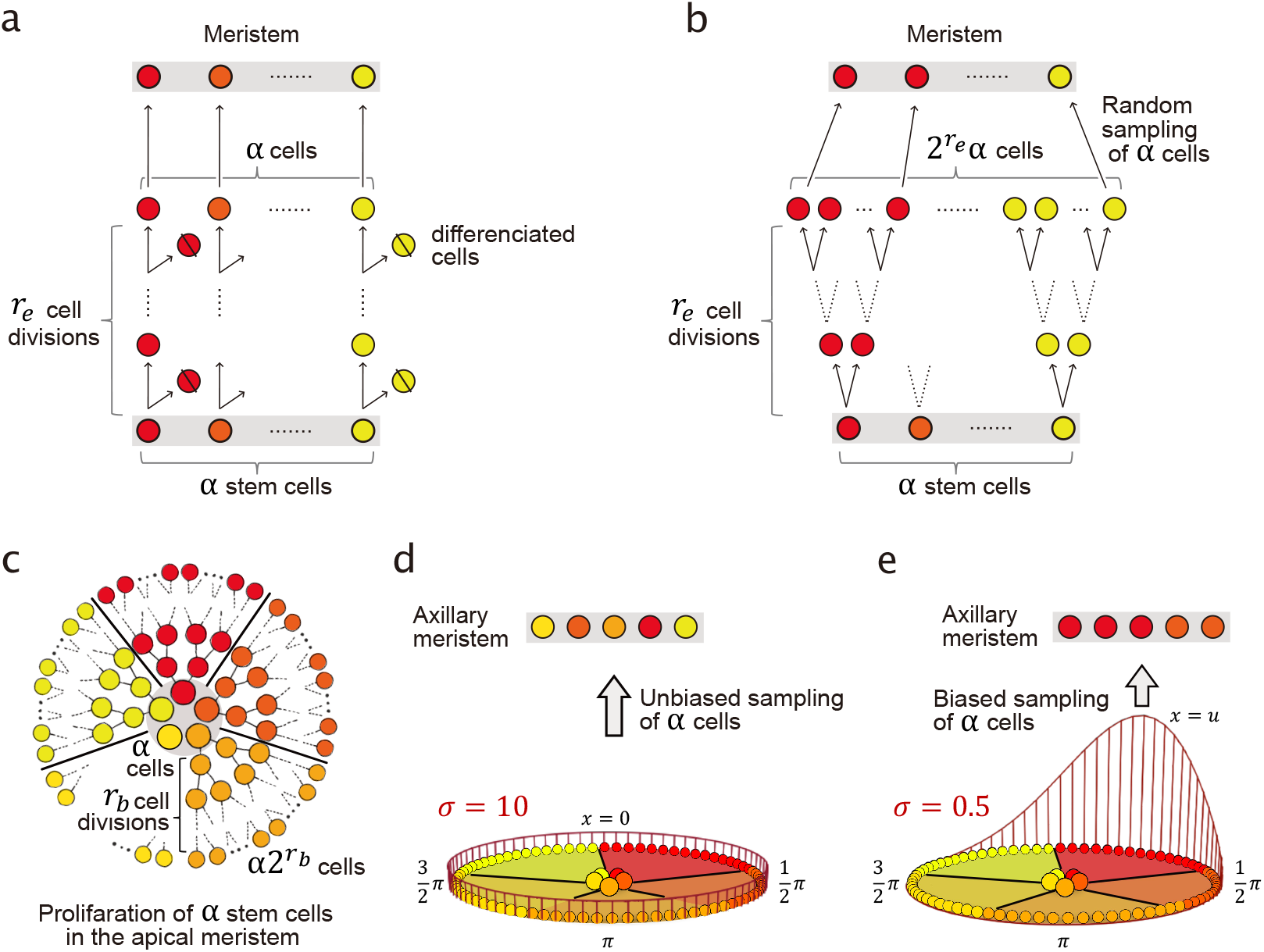
Schematic representation of the models of elongation (a, b) and branching (c-e). Each circle represents a cell, and stem cell initials are highlighted by the grey background (*α* = 5). Stem cell lineages are denoted by the different colors. (a) The structured elongation. During *r_e_* cell divisions, one side of the daughter cells are eliminated and differentiated into tissue (denoted by the slashed circle), and the other side of daughter cells are deterministically to be subsequent initials without sampling. Stem cell lineages are maintained during elongation. (b) The stochastic elongation. None of the daughter cells are eliminated, and from the remaining *α2^r_e_^* cells, *α* cells are randomly sampled for the next initials. (c) In branching, firstly *α2^r_b_^* cells are generated through *r_b_* cell divisions of the stem cells in the apical meristem (on the centre grey circle). (d, e) Secondly, from *α2^r_b_^* cells, cells for axillary meristem are sampled with weights denoted by the red line. (d) The unbiased sampling (*σ* = 10). Cells are sampled completely at random. (e) The biased sampling (*σ* = 0.5). Cells around *x* = *u* are more likely to be sampled. Mutations, omitted in the figures, stochastically occur during cell divisions for each daughter cell during elongation and branching.

In contrast, stochastic meristems have a population of stem cells in which constant shifting and changing of initial cells occur. The stochastic meristem can represent apices of gymnosperms with a weak tunica-corpus distinction (Klekowski et al., 1985). The stochastic meristem is modelled by random sampling of *α* stem cells from the pool of daughter cells generated by the successive divisions from *α* stem cells during elongation. When the number of cell divisions during elongation per unit length is *r_e_*, the size of the daughter cell pool becomes *α2^r_e_^* (Fig. 2b). While the structured meristem is modelled assuming the maintenance of spatially separated cell lineages, the stochastic meristem corresponds to the case where spatial structure in the meristem is lost due to the disorderly cell divisions, e.g., periclinal division.

### 3.2. Modelling branching

Branching is the process in which the axillary meristem is generated from cells arising from the apical meristem. We modelled this process by assuming that *α* stem cell initials proliferate radially by the successive cell divisions of *r_b_* times (Fig. 2c), and the *α* stem cell initials for the newly formed axillary meristem are sampled from *α2^r_b_^* cells (Fig. 2d, e).

In the sampling process of stem cells for the newly formed axillary meristem, *α2^r_b_^* cells are arranged on the circumference of the unit circle. Cells generated from the same mother cells are positioned closer than others according to their cell lineages (Fig. 2c). Let *x* ∈ [0,2π) be the position on the circumference, where the cells are arranged. The probability that the cell that occupies position *x* is sampled to form a new axillary meristem is expressed as follows: *f*(*x*; *u*, *σ*) R, where *f*(*x*; *u*, *σ*) is the probability of a wrapped normal distribution, and *R* = 2*π*/*α*2^*k*^ is the standardized constant indicating the area occupied by a single cell. The probability density function of the wrapped normal distribution is given as follows:

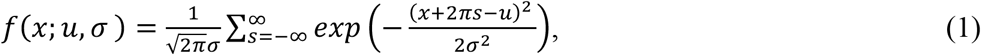

where *u* ∈ [0,2π) and *σ* > 0 are the mean and standard deviation of the distribution, respectively. If *σ* is large enough, the distribution is close to the uniform distribution, and cells for the axillary meristem are sampled completely at random (unbiased sampling; Fig. 2d). On the other hand, if *σ* is small, the cells that occupy the position around *x* = *u* are more likely to be chosen, resulting in biased sampling of a particular stem cell lineage (biased sampling; Fig. 2e). Thus, cells in the vicinity of a place where axillary meristem will be formed are apt to contribute to the formation of the axillary meristem. The former unbiased sampling corresponds to the case where proliferating cells expand with a high migration rate across the boundaries between cell lineages, resulting in the axillary meristem formation under the well-mixed cell lineages. In contrast, the latter biased sampling corresponds to the case where cell lineages are stable with strict boundaries for each cell lineage. For each branching process, *u* is randomly sampled from circumference, [0,2*π*). To perform numerical simulation of the model, we approximated Eq. (1) according to Kurz et al. (2014).

### 3.3. Modelling mutation

Somatic mutation is assumed to occur in each stem cell under cell division during elongation and branching processes at a rate *v* per cell division. The mutation occurs in each of the two daughter cells independently, as assumed in a previous study (Klekowski et al., 1989). When mutation occurs in a stem cell, the mutated genomic site is randomly chosen from the whole genome of size *G* bp. Thus, the mutation rate per cell division per genomic site (*μ*) is equal to *v/G*. Given the sufficiently small mutation rate *μ*, back mutations are negligible.

The state of the *k*-th site (*k* ∈ [1,6]) of the *i*-th stem cell (*i* ∈ [1,*α*]) in the meristem of branch *n* can be either nonmutated (0) or mutated (1) and is formalized as follows:

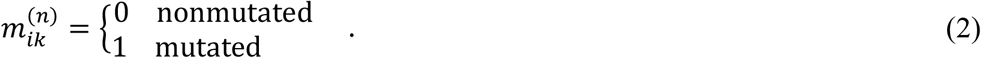

We define the stem cell state by Eq. (2), and the meristem state of genomic site *k* at a branch *n* is given as 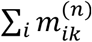, the number of mutated cells in the meristem (Fig. 3). In addition, branch *n* is defined as mutated at site *k*, if 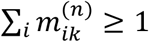, the mutated cell is present in the meristem. Then, the number of mutated sites in each branch is defined as the number of sites that satisfy this. In addition, fixation of the somatic mutation at branch *n* is also defined by 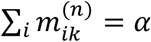, which represents the case when all stem cells in the meristem are mutated at the focal site *k* (Fig. 3). Since no empirical studies to date have provided evidence of natural selection against somatic mutations within trees (Orr et al., 2019; Perez-Roman et al., 2021; Duan et al., 2022; Imai et al., 2023), we focused exclusively on the neutral mutations in our model.

**Figure 3.**
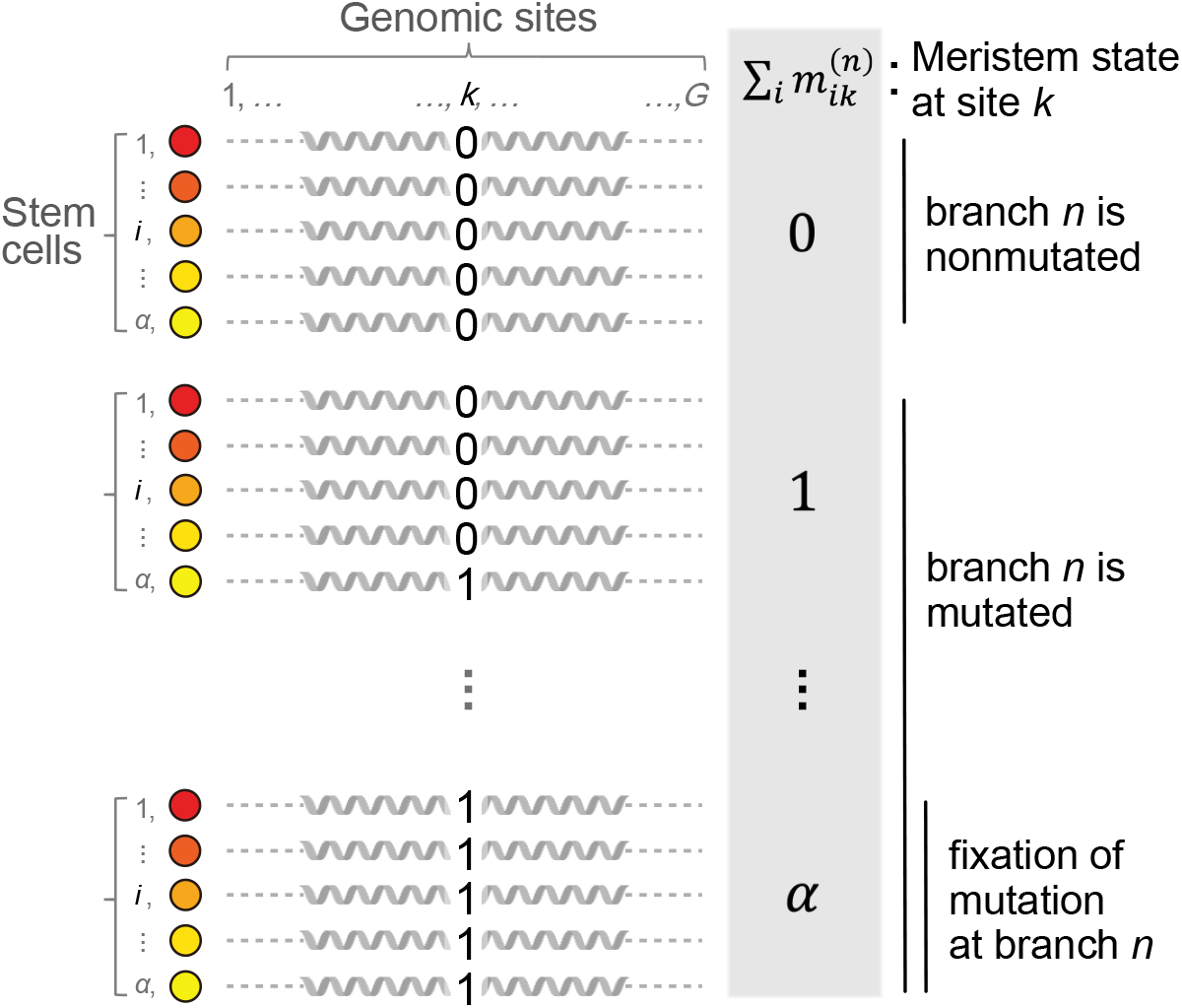
Schematic representation of the meristem state of genomic site *k*.

### 3.4. Four models with different elongation and branching processes

We analysed four extreme models with varying elongation and branching processes (Table. 1) and examined how each process affects the accumulated number and the distribution of somatic mutations across branches. With regard to the elongation process, we focused on structured and stochastic meristems. During elongation of the structured meristem, replacement of stem cell lineages never occurs (Fig. 2a), but in the stochastic meristem, replacement of stem cell lineages frequently occurs (Fig. 2b). With regard to the branching process, we examined unbiased and biased branching, namely, *σ* = 10 and 0.5, respectively. If *σ* = 10, the sampling probability given in Eq. (1) is close to the uniform distribution, and stem cells for the axillary meristem are sampled at random (Fig. 2d). However, if *σ* = 0.5, the sampling probability centred at *x* = *u* and stem cells for the axillary meristem are sampled more frequently from cells in the vicinity of *x* = *u* (Fig. 2e). Taken together, we focused on 2 × 2 models (Table. 1).

**Table 1.**
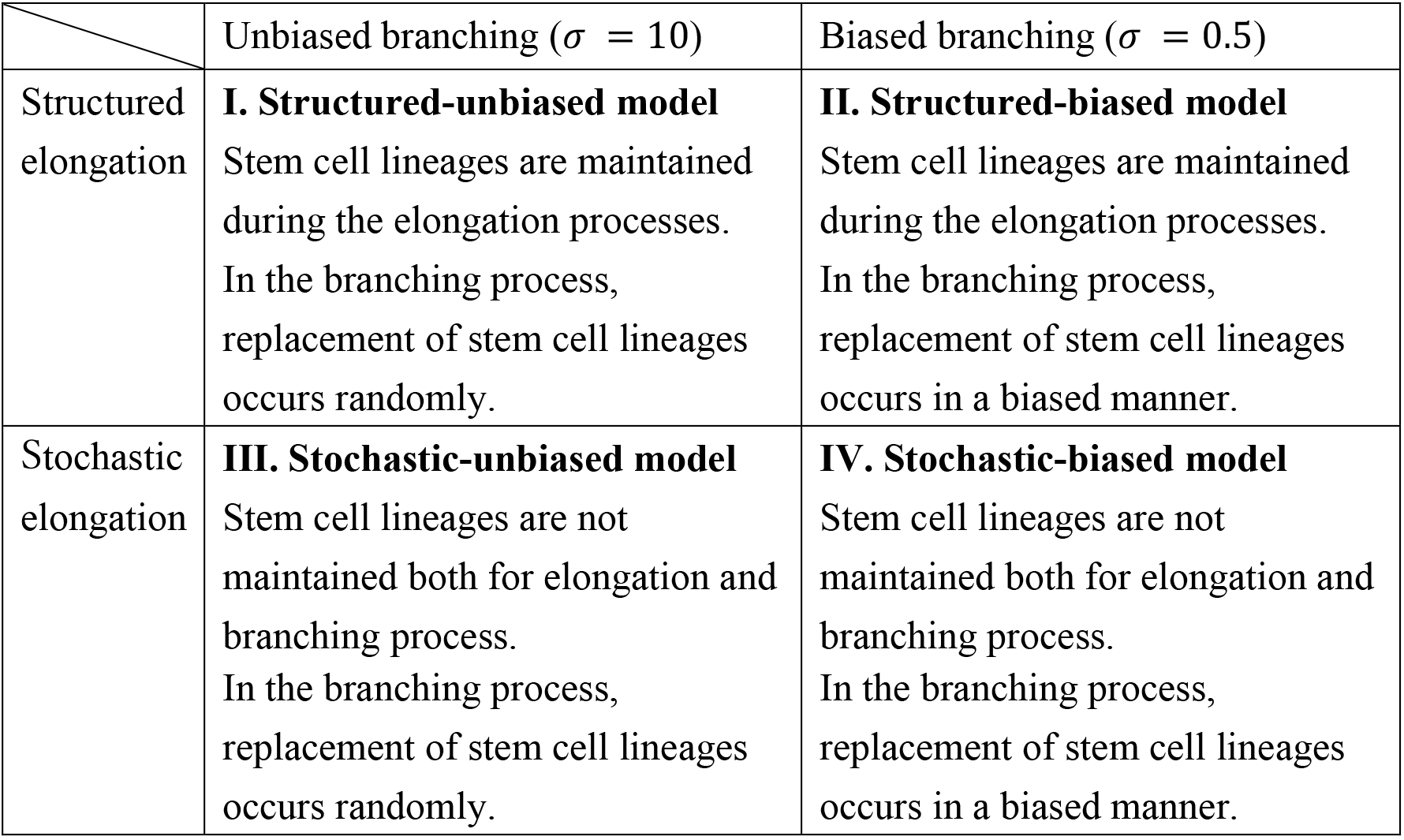
Four models with different elongation and branching processes.

### 3.5. Mathematical formulation of somatic mutation accumulation

To confirm simulation results from the elongation process in the structured and stochastic models, we used a mathematical model proposed by Klekowski et al. (1989) by focusing on the somatic mutation accumulation at a single site in a single branch. Here we briefly explain about the model. Note that, in this mathematical model, we considered a mutation at a single site, whereas mutations at multiple genomic sites are considered in the simulation model described before. In addition, we exclusively focused on the case when *r_e_* = 1, as in the application of the simulation model (Table 2). Let *i* be the number of mutated cells at an arbitrary site in a population of *α* stem cells (0 ≤ *i* ≤ *α*). We introduced the state vector ***π***(*t*), which represents the probability that the stem cell population includes *i* mutated cells at time *t*:

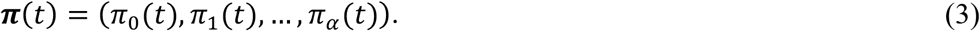

In the structured model, the number of mutated cells changes from *i* to *j* with a transition probability that is given as the binomial distribution as follows:

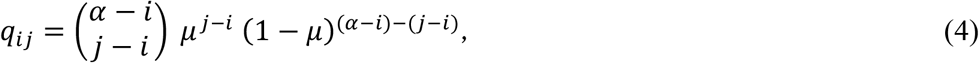

where *μ* is the mutation rate per cell division per site. Eq. (4) indicates the probability that *j-i* cells are mutated while others remain nonmutated. Note that Eq. (4) holds for *j* ≥ *i*; otherwise, *q_ij_* = 0. By introducing a transition matrix ***Q*** = [*q_ij_*] (0 ≤ *i,j* ≤ *α*)), the state change of the stem cell population in the structured model is given as follows:

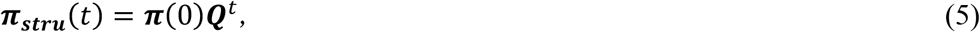

where ***π***(0) = (1,0,…, 0) is the initial condition in which there are no mutated cells.

**Table 2.** Parameters used in the simulation models

| Symbol | Definition | Values in the simulation |
| --- | --- | --- |
| $\alpha$ | Number of stem cells in the shoot meristems | 5 |
| $r_e$ | Number of cell divisions during elongation | 1 |
| $r_b$ | Number of cell divisions during branching | 7 (Burian et al., 2016) |
| $\sigma$ | Degree of sampling bias during branching | 0.5 or 10 |
| $u$ | Center position of newly formed branch | $u \sim U(0, 2\pi)$ |
| $\mu$ | Mutation rate per site per cell division | $1.33 \times 10^{-10}$ (Hofmeister et al., 2020) |
| $G$ | Genome size | 54,998,919<br>(Hofmeister et al., 2020) |
| $v$ | Mutation rate per cell per cell division (in a whole genome) | $7.31 \times 10^{-3} (\mu G)$ |

In the stochastic model, the number of mutated cells changes from *i* to *j* with a transition probability that is given as follows:

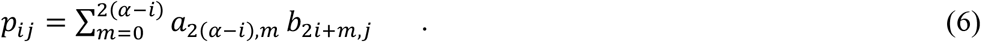

Here, *a*_2(*α-i*),*m*_ is the probability that *α* - *i* nonmutated cells generate *m* mutated cells after single cell division and is calculated as follows:

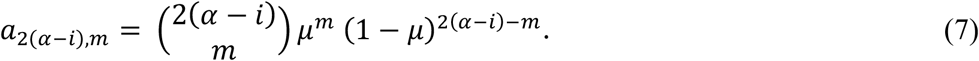

After a cell division, the number of non-mutated daughter cells becomes 2(*α* — *i*). Therefore, the probability that *m* daughter cells are selected to be mutated is given as Eq. (7). After the mutation event, the number of mutated daughter cells becomes 2*i* + *m*. The probability that *j* mutated cells are sampled from 2*i* + *m* daughter cells and *α* — *j* non-mutated cells are sampled from 2*α* — (2*i* — *m*) daughter cells is given as

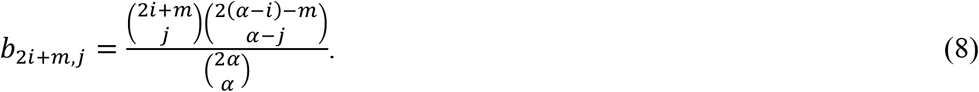

Overall, the probability that the number of mutated cells changes from *i* to *j* is given as the product for all possible cases as given in Eq. (6) (see Klekowski et al., 1989). By introducing a transition matrix ***P*** = [*p_ij_*], (0 ≤ *i,j* ≤ *α*) the state change of the stem cell population in the structured model is given as follows:

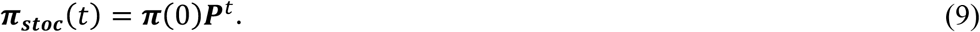

Based on Eqs. (5) and (9), the expected number of mutated cells at an arbitrary site for the structured and stochastic model are represented by ∑_*i*_ *i π_stru_i__*(*t*) and ∑_*i*_ *i π _stoc_0__* (*t*), respectively. At the meristem level, the probabilities that the meristem includes at least one mutated cell in the structured and stochastic models are given by 1 — *π_stru_0__*(*t*) and 1 — *π*_*stoc_0_*_ (*t*), respectively. Note that the mathematical models accurately predicted the simulation results (Fig. A1c–d).

## 4. Application of the model to the 330-year-old popular tree

### 4.1. Simulation of mutation accumulation in a poplar tree with hierarchical modular architecture

To simulate mutation accumulation and expansion in silico, we applied our models to a 330-year-old *P.trichocarpa* tree in which branch-level genotyping and age estimation of each branch were performed previously (Fig.4a; Table A1, A2; Hofmeister et al., 2020). Based on genome sequencing of leaves from eight branches in a single individual, Hofmeister et al. (2020) detected single nucleotide polymorphisms (SNPs) among different branches and estimated the mutation rate per year per site using the estimated age of each branch. Because the number of cell divisions per year is not known, we assumed that the number of cell divisions during elongation is once per year to grow the unit length (*r_e_* = 1). Based on this assumption, we converted the estimated mutation rate per year per site (1.33×10^-10^) to per cell division, dividing by *r_e_* (*μ* =1.33×10^-10^/*r_e_*). Then, we obtained the mutation rate per cell division per cell *v* = *μG* = 7.31×10^-3^, by multiplying the genome size used for SNP detection (*G* = 54,998,919; values given in Table S3 in Hofmeister et al. 2020) to *μ*. We applied our model to the observed modular architecture of the popular tree, which has two main axes, branches 4 and 8, with three lateral branches per each axis (Fig.4a; Table. A1, A2). We assumed that the size of the stem cell population is five (i.e., *α* = 5) and at the beginning of growth none of them possesses any mutations as the initial condition. According to the empirical data from *A. thaliana* and tomato (Burian et al., 2016), we assumed that stem cell initials of a newly formed axillary meristem experience seven cell divisions (i.e., *r_b_* = 7). Mutations in the branching process also occur at the same rate *v*. All the parameter values used in the simulation are listed in Table 2.

**Figure 4.**
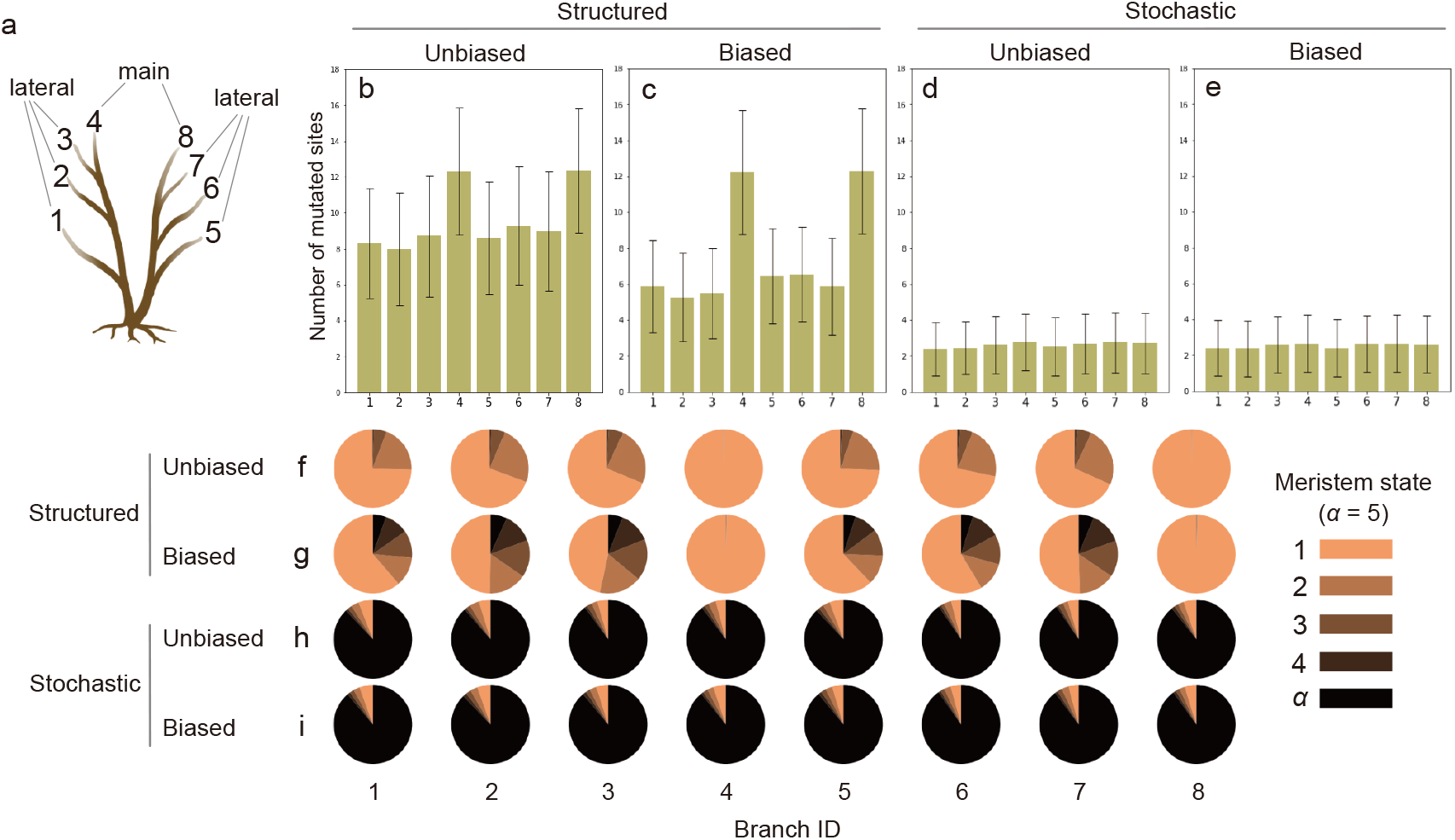
The accumulated mutations at the tip of branches. (a) The modular architecture of the poplar tree (from Hofmeister et al. 2020). Numbers indicate branch IDs, where branches 4 and 8 are the main axes, and branches 1–3 and 5–7 are lateral branches. (b-e) Mutations accumulated in the meristems at the tip of each branch. Each bar represents the number of mutated sites in each branch, and the error bars represent the standard deviation. (b) The structured-unbiased model. (c) The structured-biased model. (d) The stochastic-unbiased model. (e) The stochastic-biased model. (f-i) The frequency of mutations for each meristem state at the tip of each branch. Colours show the state of mutations; lighter colour indicates chimeric mutations which are shared by a part of stem cells, and black indicates the fixed mutations. (f) The structured-unbiased model. (g) The structured-biased model. (h) The stochastic-unbiased model. (i) The stochastic-biased model. The results in (b-i) are the average of 1000 simulations.

We counted predicted mutated sites in each branch and compared these values with the observed SNP counts for each branch of the poplar tree (Hofmeister et al., 2020). Given that SNP data are derived from the sequence of bulk samples that include many cells, cell-level mosaicism cannot be evaluated in these data. When we compare model predictions with the data, we considered that the branch *n* is mutated at site *k* when the mutated cells are present in the meristem, namely, 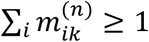 based on Eq. (2). This assumption indicates that rare mutations in a stem cell population can be detected. We performed 1000 independent simulations and calculated the average value to compare predictions with observations.

### 4.2. Mutation distribution on a modular architecture of the poplar tree

The four different models generate different consequences of mutation expansion in a modular architecture. To evaluate the mutation expansion patterns, we examined the distribution of focal mutated sites over eight branches. We introduced a vector with eight columns, corresponding to the presence (1) or absence (0) of the mutated site in branch *n* (1 ≤ *n* ≤ 8). For example, [1,0,0,0,0,0,0,0] indicates that the mutation at a focal site is present only at branch 1. Given that eight branches were considered for this analysis, there are 2^8^ - 1 possible distribution patterns of a mutation after subtracting a single pattern corresponding [0,0,0,0,0,0,0,0], i.e., without mutation. For each mutated site, we classified the distribution pattern across branches and drew the histogram for the number of cases for each distribution pattern.

To compare predicted and observed patterns of somatic mutation distribution, we calculated the mean squared error (*MSE*) that measures the amount of error in the model as follows:

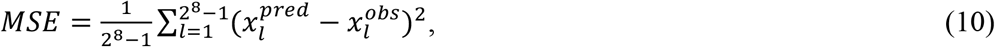

where 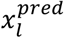 and 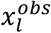 represent the predicted and observed number of mutated sites distributed in the *l*-th pattern, respectively. There are 2^8^ — 1 possible patterns of mutation distribution. We performed 250 independent simulations, and the *MSE* averaged over each simulation was calculated.

## 5. Results

### 5.1. Comparison of chimeric patterns within a stem cell population at the branch level

The number of mutated sites in a meristem, which is defined as the number of genomic sites that includes at least one mutated cell in a stem cell population, varied among models. In each of eight branches, the number of mutated sites was predicted to be greater in the structured models compared with stochastic models (Fig.4b-e). This finding is attributed to the loss of mutation during the sampling process of stem cells in the stochastic models, i.e., somatic genetic drift. This result is confirmed by the mathematical model formalized in Eqs. (5) and (9) by exclusively considering the elongation process (Fig. A1). The mathematical model showed that the probability that mutation is present in at least one stem cell in a meristem increases almost linearly with branch age in both the structured and stochastic models. In a young branch, however, the increasing trend becomes nonlinear in the stochastic model (Fig. A1a). Note that despite the difference in the probability of the presence of the mutated cell between the structured and stochastic models, the expected number of mutated cells calculated from Eqs. (5) and (9) were equivalent between the two models. The branching process is another driver for the loss of mutations in lateral branches. The meristem in the lateral branches (branches 1-3 and 5-7) possessed fewer mutated sites than in the main axis (branches 4 and 8; Fig.4a) in the structured model (Fig. A2a, b). Moreover, the number of mutated sites in lateral branches was fewer in the structured-biased model compared with the structured-unbiased model (Fig.4b, c; Fig. A2a, b), due to the loss of mutations during the biased formation of the axillary meristem. In the stochastic models, no differences in the number of mutated sites were noted between the main axis and lateral branches (Fig. A2c, d) due to the fixation of most mutated sites in the stem cell population of the meristem, during the elongation process prior to the branching process.

Somatic mutations are not always fixed; rather, somatic chimaera can be maintained in a stem cell population for a long-time during the growth and survival of a tree. To evaluate each model in the light of chimerism, a somatic mutation at the focal site is characterized by *α* different states, depending on the number of mutated cells in the meristem (Fig. 3). For example, the meristem state is *α*, if the mutation is fixed in the stem cell population. The frequency of somatic mutations for each meristem state differed among models, especially between structured and stochastic models (Fig.4f-i). In the structured models, the frequency of the state-1 mutation was dominant, indicating that most mutations exist only in a single cell in a meristem characterized by a somatic chimaera. The frequency from the state-2 to state-*α* mutation was higher in the structured-biased model compared with the structured-unbiased model because the biased sampling of stem cells in the branching process increases the frequency of mutation fixation (Fig.4f, g). At the main stems (branches 4 and 8), almost all mutations were characterized by the state-1 meristem because cell lineages are maintained throughout elongation (Fig. A1b). In contrast, the state-*α* mutation was dominant in the stochastic models due to the fixation of mutations by somatic genetic drift during elongation. The frequency of each state was unchanged between stochastic-unbiased and -biased models and among the main and lateral branches (Fig.4d, e, h, i). These results suggest that the sampling effect during branching on mutation accumulation was negligible compared to the sampling effect during elongation.

These results demonstrate that the degree of intra-meristem mosaicism is higher in the structured than in the stochastic models. In addition to intra-meristem mosaicism, inter-meristem mosaicism was assessed by calculating variance in the frequency of mutated cells in the meristem across branches. Here, the frequency of mutated cells at genomic site *k* at branch *n* is given as based on Eq. (2). As expected, the variance of the frequency across branches averaged over all mutated sites was larger in the stochastic models (0.146 ± 0.0315 and 0.147 ± 0.0307 for the stochastic-unbiased and -biased models, respectively) compared with the structured models (0.0125 ± 0.00255 and 0.0221 ± 0.00595 for the structured-unbiased and -biased models, respectively). Greater variation in the frequency of mutated stem cells across meristems highlights the higher degree of inter-meristem mosaicism in the stochastic models. We also found that the variance in the biased models was greater than that in the unbiased models, suggesting that biased sampling of stem cells during branching also contributes to somatic mosaicism at the individual level.

### 5.2. Somatic mutation distributions across branches in a tree

The most frequently observed mutated pattern was singleton, in which the mutation was present only at a single branch (Fig. 5). All eight possible distribution patterns of singletons were present in all four models, which was consistent with the data from the poplar tree. Twin mutations in which the mutation was present at two branches were less frequent than singleton mutations (Fig. A3). Among the predicted twin mutation, the mutations that reflect tree topology were dominant. For example, the twin mutation [0,0,1,1,0,0,0,0], in which a new mutation emerged before budding of branch 3 and expanded to both branches 3 and 4, was frequently observed in all models. In contrast, the mutations that do not reflect tree topology were predicted exclusively in structured models. For example, the twin mutation, [0,1,0,1,0,0,0,0], in which a new mutation expanded to branches 2 and 4 but was lost in branch 3, was predicted only in the structured models (Fig. 5a, b). The observed twin mutations in the poplar tree showed the presence of topology-independent twin mutations, although the count was very small. Two types of twin mutations were present in the data but were absent in model predictions (Fig. 5d). These twin mutations were [0,1,1,0,0,0,0,0] and [0,0,0,0,0,1,1,0], in which mutations were present only in lateral branches.

**Figure 5.**
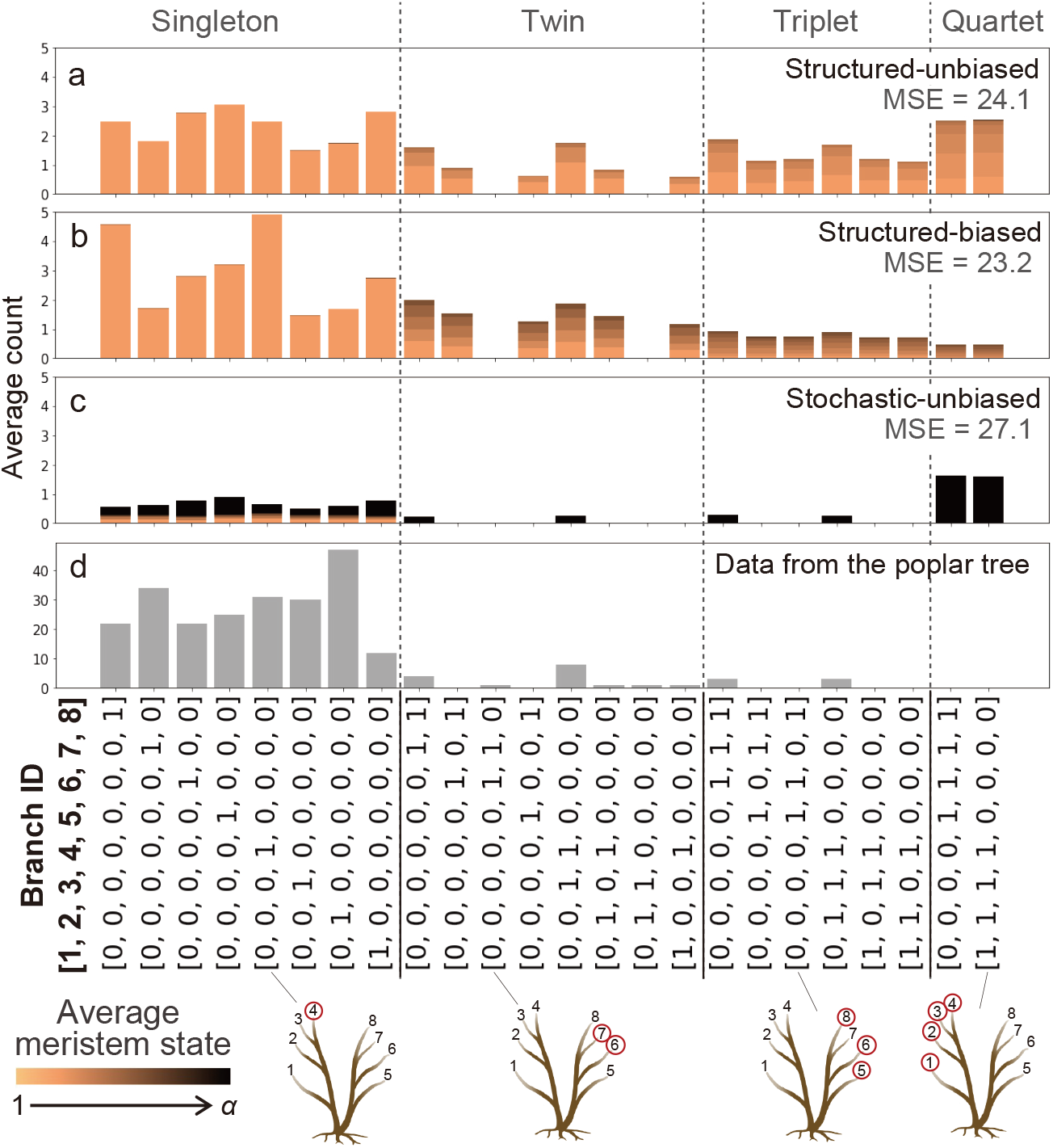
The distribution of somatic mutations within the modular architecture. Each bar represents the average number of mutated sites distributed in a pattern. Patterns under 0.1 are omitted here. Colours show the averaged meristem state of a mutation over focal branches. (a) The structured-unbiased model. (b) The structured-biased model. (c) The stochastic-unbiased model. The stochastic-biased model is omitted here because the result was very similar to the stochastic-unbiased model. The results in (a-c) are the average of 1000 simulations. (d) The data from the sequenced poplar tree listed in Tables S1 and S2 of Hofmeister et al. (2020). RR (homozygous) and RA (heterozygous) in the tables were regarded as 0 (nonmutated) and 1 (mutated), respectively, because all detected mutations in the poplar tree are from homozygous to heterozygous, according to Hofmeister et al. (2020). The *MSE* from the poplar tree was 24.1, 23.2, 27.1, and 27.1 in the structured-unbiased, structured-biased, stochastic-unbiased, and stochastic-biased models, respectively. When quartets were excluded from the calculation, the MSEs were 23.9, 23.3, 27.0, and 27.0 in the same respective order.

With regard to the triplet mutations, topology-independent mutations, such as [1,1,0,1,0,0,0,0] and [0,0,0,0,1,0,1,1], were also observed in the structured models but not in the stochastic models or in the data. We also found a striking difference between the data and model predictions. Specifically, all models predicted the presence of quartet mutations in which the mutation was fixed in four branches (branches 1–4 and branches 5–8), but such quartet mutations were not observed in the poplar tree (Fig. 5d). This finding is likely attribute to the SNP detection method used in the previous study, as we discussed later. Overall, the observed mutation distribution patterns were most similar to those of the structured-biased model: the *MSE* from the poplar tree was the smallest in the structured-biased model (Fig. 5). Even if quartets are excluded in the calculation, the trend does not change (see the caption of Fig. 5).

The predicted distributions of somatic mutations across branches were not significantly different when we increased the number of stem cells in the meristem (*α*) from five to 15 (Fig. A4); however, increased *α* resulted in an increase in somatic mutations and a decrease in fixed mutations (Fig. A1), as predicted in previous theoretical studies (Klekowski and Kazarinova-Fukshansky, 1984a, b; Klekowski et al., 1989; Pineda-Krch and Fagerström, 1999).

## 6. Discussion

We simulated the accumulation and expansion of somatic mutations in the modular architecture of the real poplar tree and compared the prediction of the models with the sequenced data. By extending the previous models, we demonstrated that the strong somatic genetic drift decreases intra-meristem mosaicism but increases inter-meristem mosaicism. Our model also revealed that the loss of mutation due to somatic genetic drift during branching leads to the mutation accumulation pattern that is different from the tree topology.

The strong somatic genetic drift during elongation in the stochastic meristem results in low intra-meristem but high inter-meristem mosaicism. The high inter-meristem mosaicism suggests the genetically distinct branches and less integrity within an individual tree. In such trees, genetically diverse offspring can be produced even by self pollination because stigma can be fertilized by pollens from different branches in the same individual. The increased genetic variation in the offspring allows natural selection to operate and increases the evolutional potential. The increased availability of somatic mutation data in the future will enable us to demonstrate the relationship between the strength of somatic genetic drift and the provision of genetic variations in the offspring population.

The somatic genetic drift also occurs during branching, resulting in the loss and fixation of somatic mutations (Klekowski et al., 1989; Burian et al., 2016). In this study, we showed that the biased sampling of cells to form a new branch enhances the effect of somatic genetic drift (Fig. A2a, b; Fig.4f, g). The effects of somatic genetic drift during branching were considerably greater in the structured models compared with the stochastic models because intra-meristem mosaicism was maintained during structured elongation in the apical meristem prior to branching. Although the contribution of branching was not as explicit in the present poplar tree with the simple architecture, branching is expected to have a greater impact within trees with a more complex modular architecture, especially in the species with structured meristem.

We also found that the mutation distribution pattern does not necessarily reflect tree topology. Some mutations that did not follow the tree topology were predicted in the structured models and observed in the poplar tree (Fig. 5). Some previous empirical studies preferentially filtered out candidate mutations not following tree topology (Wang et al., 2019; Orr et al., 2020). Our results suggest that such filtering would underestimate the accumulation of somatic mutations. Some discrepancies in the distribution pattern were noted between the prediction and the observation. First, in contrast to the predictions, no quartet mutation was observed in the data from the poplar tree (Fig. 5d). This finding may be attributed to the SNP detection method, and samples from eight branches were not pooled (Hofmeister et al. 2020). Therefore, mutations that segregate the genotypes between modules (branches 1–4 and branches 5–8) could not be detected. Next, the two patterns, [0,1,1,0,0,0,0,0] and [0,0,0,0,0,1,1,0], were not predicted in the models but observed in the poplar tree. These differences are explained by the possibility that branches 4 and 8 were not the main stem but were lateral branches. In this case, these patterns are predictable. In addition, the counts of mutations were much higher in the data from the poplar tree compared with the prediction (Fig. 5). This finding may be because, in the data used to plot the mutation distribution, mutated sites out of effective genome size that were not used in the estimation of mutation rate were included. Given that false positive mutations were reported as singletons and that the number of singletons was excessive in the data (Fig. A3e), errors were possibly included in these mutated sites. In addition, the much larger size of the stem cell population *α* in the poplar tree is also considered. At least in this study, when *α* = 5, the structured-biased model predicted the closest pattern to the observation in the poplar tree. This result may suggest that the branching process, rather than elongation, was the main driver of somatic genetic drift in this tree. Since the stable stem cell lineages in the structured model are assumed to be the meristem of angiosperm and the poplar tree is angiosperm, this result was in line with our assumptions. However, more comprehensive analyses based on high-quality somatic mutation data are necessary to conclude. In the near future, with technological progress and the accumulation of high-quality data, it will be possible to perform more robust parameter estimation through model fitting, which is an intriguing subject for future work.

In the structured model assumed to correspond to the angiosperm, most of the mutations existed exclusively in a few stem cells in a meristem (Fig.4f, g; Fig. A1). Hence, as Plomion et al. (2018) and Reusch et al. (2021) noted, some mutations were possibly overlooked in these studies. In a recent study, intra-meristem genetic mosaicisms were estimated by ultradeep resequencing (Yu et al., 2020). Single-cell DNA sequencing also has the potential to unveil mosaicism within the meristem (Reusch et al., 2021). In addition, the methodology may enable us to estimate the per cell somatic mutation rate independent of somatic genetic drift during elongation. Because the expected number of mutated cells calculated from Eqs. (5) and (9) were equivalent, the number of mutations per cell, not per meristem, is constant regardless of whether structured or stochastic elongation occurs.

Taken together, our model enabled us to evaluate the effect of cell lineage dynamics on elongation and branching and showed that the accumulation and expansion patterns of somatic mutations are highly variable depending on the growth processes even under the same modular architecture and mutation rate. Moreover, by comparing the prediction from the models with the observation in the real poplar tree, we estimated the behaviour of somatic mutations associated with the cell lineage dynamics invisible from the snapshot of the sequenced data, which makes it possible to understand the observed data more deeply.

## Acknowledgement

This study was funded by JSPS KAKENHI (JP17H06478) to A.S.

## A Appendix

**Figure A1.**
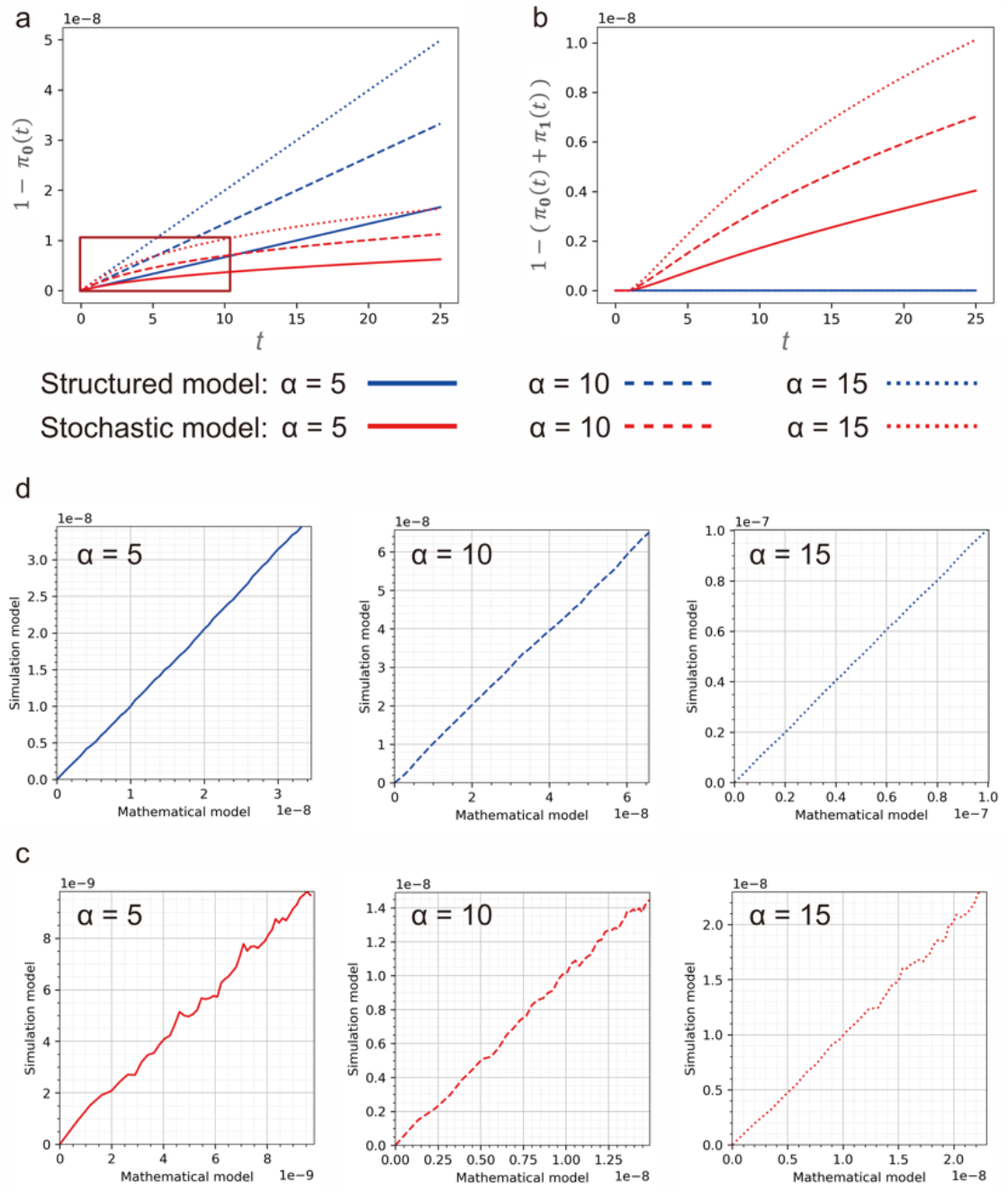
The Mathematical formulation of somatic mutation accumulation during elongation. (a) The probability that at least one stem cell is mutated in a meristem. In the vicinity of *t* = 0, which is early stage of the development, the probability of the structured 1 — *π*_stru_i__(*t*) and the stochastic 1 —*π*_stoc_i__ (*t*) overlap. However, as *t* increases, the probability of the stochastic model decreases from the structured model nonlinearly. The region of non-linear increase is highlighted by a square. (b) The probability that more than one stem cell is mutated in a meristem. Comparison with (a) shows that almost all mutations are state-1 in the structured meristem, whereas states greater than 1 are dominant in the stochastic model. (c–d) Comparing predictions from the mathematical model to computer simulations. (c) The structured model. (d) The stochastic model. The comparisons are based on the probability that the meristem includes at least one mutated cell at the focal site, and the probabilities from *t* = 0 to 50 are plotted. In the mathematical model, the probability that corresponds to 1 — *π*_0_ (*t*) was calculated using Eq. (5) for the structured model and Eq. (9) for the stochastic model. In the computer simulation, we performed 1,000 independent simulations and plotted the average. All calculations were based on the parameters listed in Table2, i.e., *μ* = 1.33 × 10^-10^ and *G* = 54,998,919.

**Figure A2.**
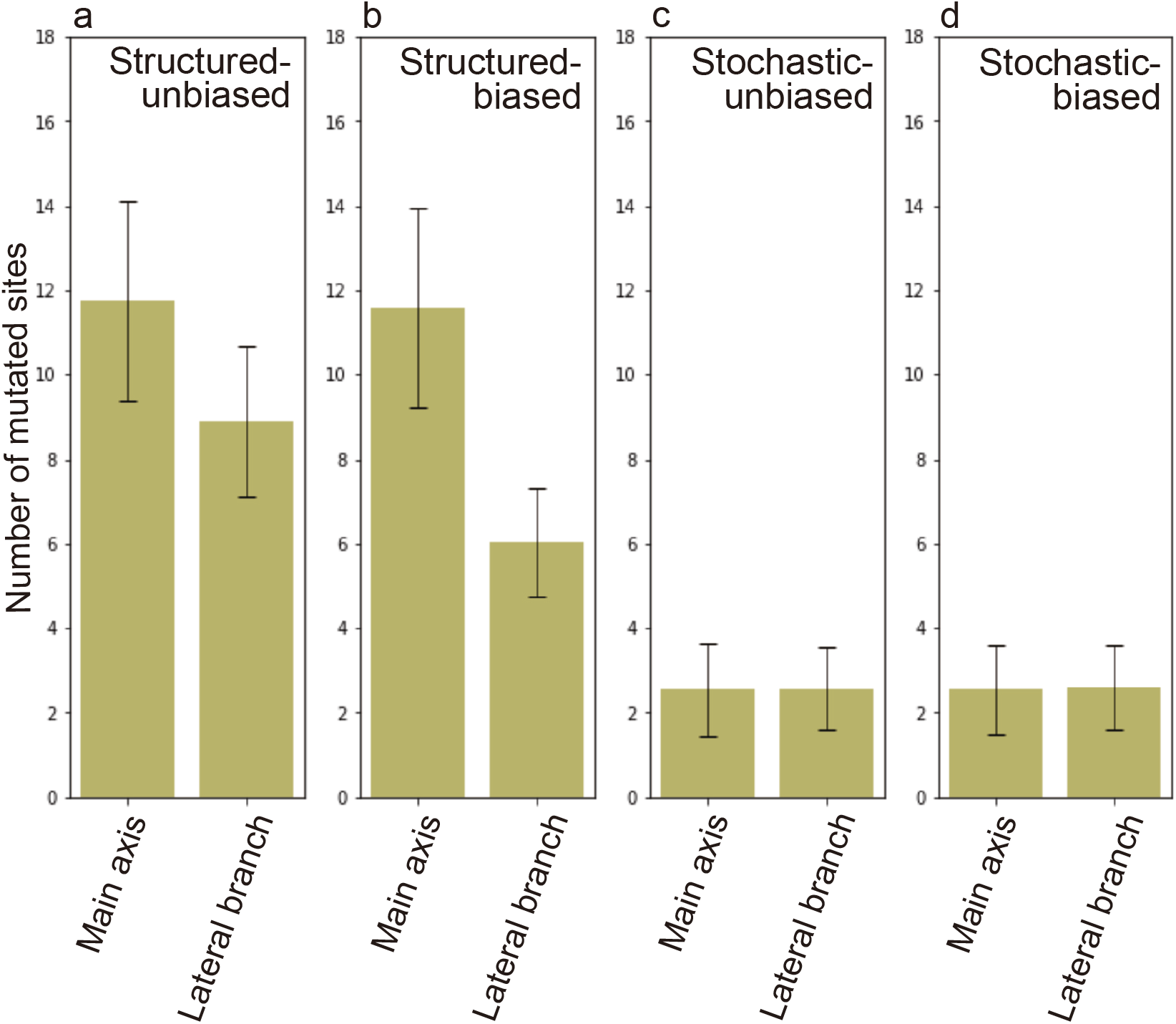
Comparison of the average number of mutated sites between the main axis (branches 4 and 8) and lateral branches (branches 1-3 and 5-7). Error bars represent standard deviation. (a) The structured-unbiased model. (b) The structured-biased model. (c) The stochastic-unbiased model. (d) The stochastic-biased model. The results in (a-c) are the average of 1000 simulations.

**Figure A3.**
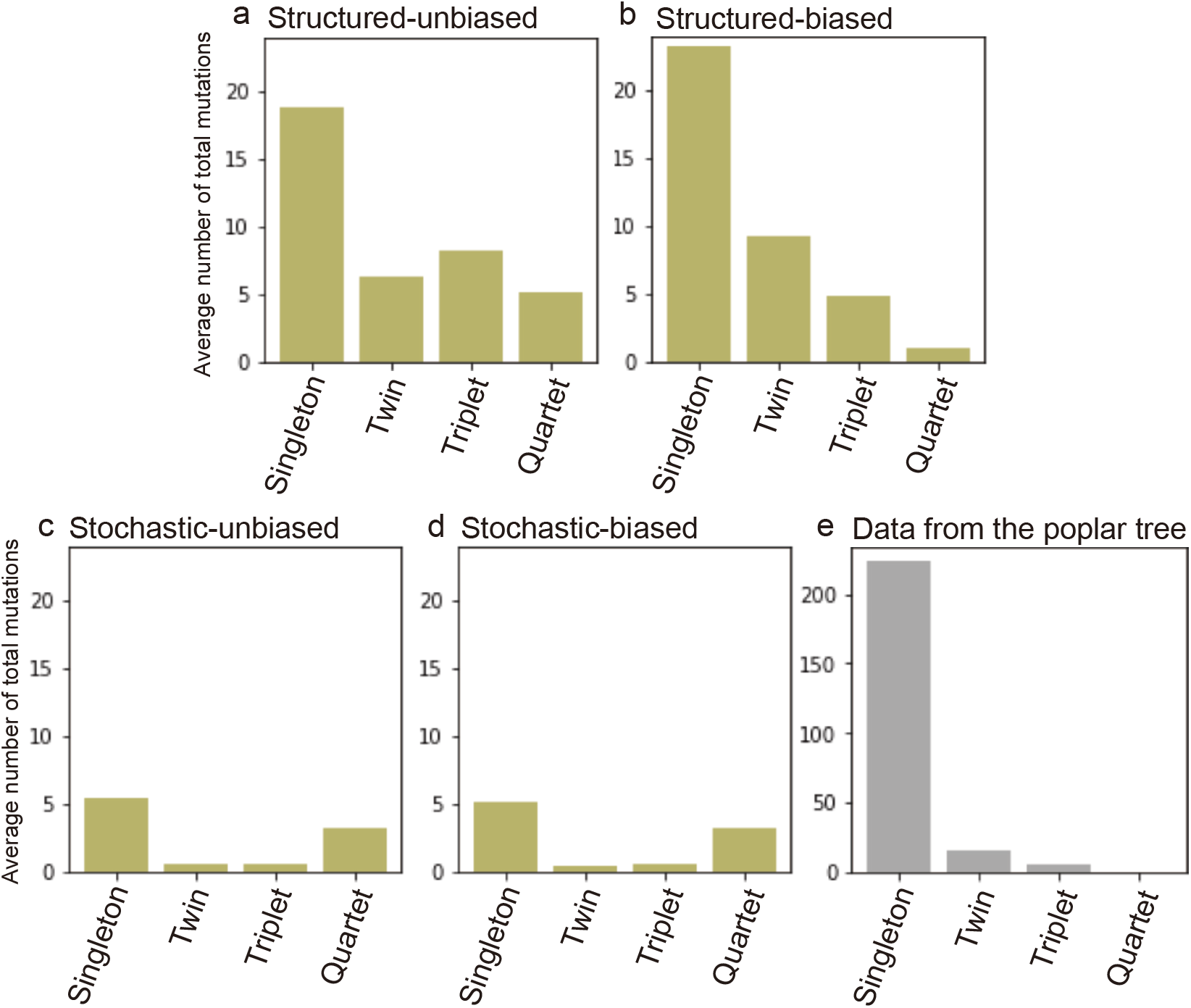
Total number of mutations summarized by the number of distributed branches. From Fig. 5, mutations distributed to the same number of branches are summarized. (a) The structured-unbiased model. (b) The structured-biased model. (c) The stochastic-unbiased model. (d) The stochastic-biased model. The results in (a-c) are the average of 1000 simulations. (d) The data from the sequenced poplar tree.

**Figure A4.**
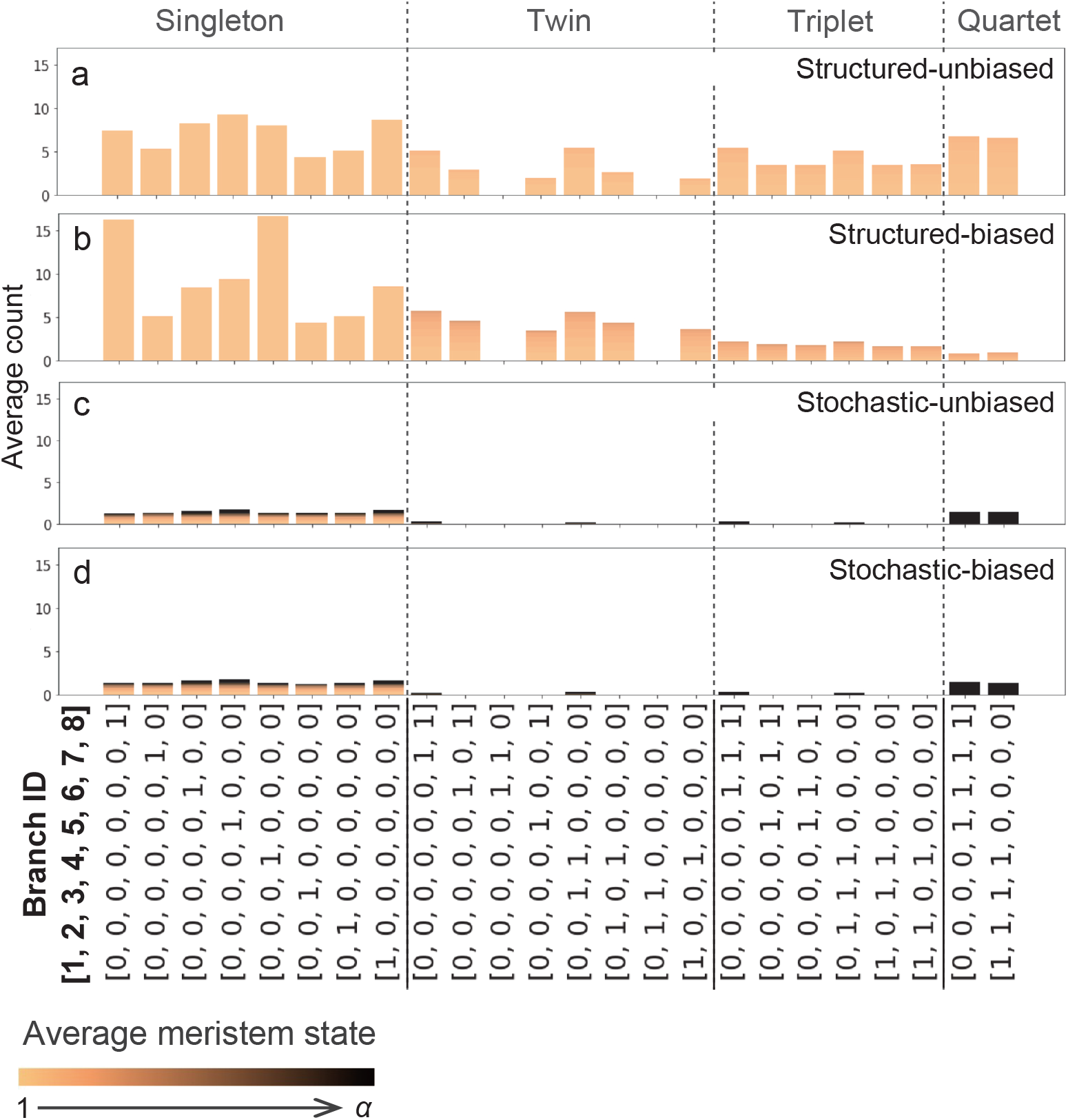
The distribution of somatic mutations within the modular architecture when *α* = 15. (a) The structured-unbiased model. (b) The structured-biased model. (c) The stochastic-unbiased model. (d) The stochastic-biased model.

**Table A1.** Estimated age of meristems at the tip of eight branches. Branches 1–4 and 5– 8 in this study correspond to branches 14.5, 14.4, 14.3, and 14.2 and 13.5, 13.3, 13.2, and 13 1 respectively in Hofmeister et al (2020)

| Branch ID | 1 | 2 | 3 | 4 | 5 | 6 | 7 | 8 |
| --- | --- | --- | --- | --- | --- | --- | --- | --- |
| Age | 289 | 290 | 320 | 332 | 297 | 328 | 327 | 330 |

**Table A2.** Estimated divergence time for each pair of eight branches

|  | 1 | 2 | 3 | 4 | 5 | 6 | 7 | 8 |
| --- | --- | --- | --- | --- | --- | --- | --- | --- |
| 1 | 0 | - | - | - | - | - | - | - |
| 2 | 145 | 0 | - | - | - | - | - | - |
| 3 | 175 | 112 | 0 | - | - | - | - | - |
| 4 | 187 | 124 | 82 | 0 | - | - | - | - |
| 5 | 582 | 583 | 613 | 625 | 0 | - | - | - |
| 6 | 613 | 614 | 644 | 656 | 191 | 0 | - | - |
| 7 | 612 | 655 | 685 | 697 | 190 | 139 | 0 | - |
| 8 | 615 | 616 | 646 | 658 | 193 | 142 | 85 | 0 |

## Reference

Antolin, M. F., Strobeck, C., 1985. The population genetics of somatic mutations in plants, Am. Nat. 126, 52–62.

Burian, A., Barbier de Reuille, P., Kuhlemeier, C., 2016. Patterns of Stem Cell Divisions Contribute to Plant Longevity, Curr. Biol. 26, 1385–1394. (https://doi.org/10.1016/j.cub.2016.03.067).

Dermen, H., 1969., Directional Cell Division in Shoot Apices, Cytologia 34, 541–558.

Duan, Y., Yan, J., Zhu, Y., Zhang, C., Tao, X., Ji, H., Zhang, M., Wang, X., Wang, L., 2022. Limited accumulation of high-frequency somatic mutations in a 1700-year-old Osmanthus fragrans tree. Tree Physiol. 42(10), 2040–2049. (https://doi.org/10.1093/treephys/tpac058).

Folse, H. J., Roughgarden, J., 2012. Direct benefits of genetic mosaicism and intraorganismal selection: Modeling coevolution between a long-lived tree and a short-lived herbivore. Evolution 66(4), 1091–1113. (https://doi.org/10.1111/j.1558-5646.2011.01500.x).

Frank, M. H., Chitwood, D. H., 2016. Plant chimeras: The good, the bad, and the ‘Bizzaria.’ Dev. Biol. 419, 41–53. (https://doi.org/10.1016/j.ydbio.2016.07.003).

Gill, D. E., Chao, L., Perkins, S. L., Wolj, J. B., 1995. Genetic mosaicism in plants and clonal animals. Ann. Rev. Ecol. Syst. 26, 423–444.

Godin, C., Caraglio, Y., 1998. A multiscale model of plant topological structures, J. Theor. Biol. 191, 1–46.

Hanlon, V. C. T., Otto, S. P., Aitken S. N., 2019. Somatic mutations substantially increase the per-generation mutation rate in the conifer *Picea sitchensis*. Evol Lett 3, 348–358. (https://doi.org/10.1002/evl3.121).

Hara, N., 1995. Developmental Anatomy of the Three-Dimensional Structure of the Vegetative Shoot Apex. J. Plant Res. 108, 115–125.

Hofmeister, B. T., Denkena, J., Colomé-Tatché, M., Shahryary, Y., Hazarika, R., Grimwood, J., et al., 2020. A genome assembly and somatic genetic and epigenetic mutation rate in a wild long-lived perennial *Populus trichocarpa*. Genome Biol. 21:259. (https://doi.org/10.1186/s13059-020-02162-5).

Imai, R., Fujino, T., Tomimoto, S., Ohta K., Na’iem, M., Indrioko, S., Widiyatno, Purnomo, S., Molla Morales, A., Nizhynska, V., Tani, N., Suyama, Y., Sasaki, E., Kasahara, M., Satake, A. 2023. The molecular clock in long-lived tropical trees is independent of growth, bioRxiv. (https://doi.org/10.1101/2023.01.26.525665).

Klekowski, E. J., 1984. Mutation load in clonal plants: a study of two fern species, Evolution 38(2), 417–426.

Klekowski, E. J., Kazarinova-Fukshansky, N., 1984a. Shoot apical meristems and mutation: fixation of selectively neutral cell genotypes. Am. J. Bot. 71, 22–27.

Klekowski, E. J., Kazarinova-Fukshansky, N., 1984b. Shoot apical meristems and mutation: selective loss of disadvantageous cell genotypes, Am. J. Bot. 71, 28–34.

Klekowski, E. J., Kazarinova-Fukshansky, N., Mohr, H., 1985. Shoot apical meristem and mutation: stratified meristems and angiosperm evolution. Am. J. Bot. 72, 1788–1800.

Klekowski, E. J., Kazarinova-Fukshansky, N., Fukshansky, L., 1989. Patterns of plant ontogeny that may influence genomic stasis. Am. J. Bot. 76, 185–195.

Klekowski, E. J., 2003. Plant clonality, mutation, diplontic selection and mutational meltdown. Biol. J. Linn. Soc. 79, 61–67. (https://doi.org/10.1046/j.1095-8312.2003.00183.x).

Kurz, G., Gilitschenski, I., Hanebeck, U. D., 2014. Efficient Evaluation of the Probability Density Function of a Wrapped Normal Distribution Sensor Data Fusion. In Proceedings of the IEEE ISIF Workshop on Sensor Data Fusion: Trends, Solutions, Applications, 1–5.

Marc, J., Hackett, W. P., 1992. Changes in the pattern of cell arrangement at the surface of the shoot apical meristem in Hedera helix L. following gibberellin treatment. Planta 186, 503–510.

Orive, M. E., 2001. Somatic Mutations in Organisms with Complex Life Histories. Theor. Popul. Biol. 59, 235–249. (https://doi.org/10.1006/tpbi.2001.1515).

Orr, A. J., Padovan, A., Kainer, D., Kulheim, C., Bromham, L., Bustos-Segura, C., et al., 2020. A phylogenomic approach reveals a low somatic mutation rate in a long-lived plant. Proc. R. Soc. B 287: 20192364. (http://dx.doi.org/10.1098/rspb.2019.2364).

Otto, S. P., Orive, M. E., 1995. Evolutionary consequences of mutation and selection within an individual. Genetics 141, 1173–1187.

Perez-Roman, E., Borredá, C., López-García, A., Talon, M., 2021. Single-nucleotide mosaicism in citrus: Estimations of somatic mutation rates and total number of variants. Plant Genome. (https://doi.org/10.1002/tpg2.20162).

Pineda-Krch, M., and T. Fagerström., 1999. On the potential for evolutionary change in meristematic cell lineages through intraorganismal selection. J. Evol. Biol. 12, 681–688.

Pineda-Krch, M., and K. Lehtilä., 2002. Cell lineage dynamics in stratified shoot apical meristems. J. Theor. Biol. 219, 495–505.

Plomion, C., Aury, J. M., Amselem, J., Leroy, T., Murat, F., Duplessis, S., et al., 2018. Oak genome reveals facets of long lifespan. Nat. Plants 4(7), 440–452. (https://doi.org/10.1038/s41477-018-0172-3).

Poething, S., 1989. Genetic mosaics and cell lineage analysis in plants. Trends Genet 5(8), 273–277.

Reusch, T. B.H., Baums, I. B., Werner, B., 2021. Evolution via somatic genetic variation in modular species. Trends Ecol. Evol. 36(12), 1083–1092. (https://doi.org/10.1016/j.tree.2021.08.011).

Satina, S., Blakeslee, A.F., Avery, A.G., 1940. Demonstration of the tree germ layers in the shoot apex of Datura by means of induced polyploidy in periclinal chimeras. Am. J. Bot. 27, 73–90.

Schmid-Siegert E., Sarkar N., Iseli C., Calderon S., Gouhier-Darimont C., Chrast J. et al. 2017. Low number of fixed somatic mutations in a long-lived oak tree. Nat. Plants 3, 926–929. (https://doi.org/10.1038/s41477-017-0066-9).

Schmidt, A., 1924. Histologische studien an phanerogamen vegetationspunkten. Bot. Arch. 8, 345–404.

Sutherland, W. J., Watkins, A. R., 1986. Somatic mutation: do plants evolve differently. Nature 320, 305. (https://doi.org/10.1038/320305a0).

Wang L., Ji Y., Hu Y., Hu H., Jia X., Jiang M., Zhang X., Zhao L., et al., 2019. The architecture of intra-organism mutation rate variation in plants. PLOS Biol. 17(4):e3000191. (https://doi.org/10.1371/journal.pbio.3000191).

Whitham, T. G., Slobodchikoff, C. N., 1981. Evolution by Individuals, Plant-Herbivore Interactions, and Mosaics of Genetic Variability: The Adaptive Significance of Somatic Mutations in Plants. Oecologia 49, 287–292.

Yu, L., Boström, C., Franzenburg, S., Bayer, T., Dagan, T., Reusch, T. B. H., 2020. Somatic genetic drift and multilevel selection in a clonal seagrass. Nat. Ecol. Evol. 4, 952–962. (https://doi.org/10.1038/s41559-020-1196-4).

Zahradníková, E., Ficek, A., Brejová, B., Vinař, T., Mičieta, K., 2020. Mosaicism in old trees and its patterns. Trees 34, 357–370. (https://doi.org/10.1007/s00468-019-01921-7).

